# BioAutoML-FAST: an automated machine-learning platform for reusable and benchmarked biological sequence models

**DOI:** 10.64898/2026.04.18.719383

**Authors:** Breno L. S. de Almeida, Robson P. Bonidia, Martin Bole, Anderson Avila-Santos, Peter F. Stadler, Ulisses N. da Rocha, André C. P. L. F. de Carvalho

## Abstract

The prediction of biological sequence properties has traditionally relied on alignment-based methods that assume evolutionary homology and depend on curated reference databases. This, in turn, limits scalability and sensitivity for large or heterogeneous datasets, remote homologs, short sequences, and rapidly evolving genomic regions. Although Machine-Learning (ML) approaches offer alignment-free alternatives, their broader adoption is limited by: (i) the lack of standardized, externally validated benchmark models across diverse datasets, and (ii) the technical expertise required for feature engineering, model selection, and evaluation. Automated machine learning (AutoML) alleviates these challenges by systematically optimizing representations and models with minimal user intervention. However, most existing frameworks prioritize task-specific model construction and lack mechanisms for preserving trained models as persistent, comparable benchmarks. We introduce BioAutoML-FAST, an end-to-end web platform for automated ML analysis of nucleotide and amino acid sequences. It supports both classification and regression tasks and automates feature extraction, model training, and evaluation without requiring prior user expertise. Uniquely, it serves as a community benchmarking resource, hosting a continuously expanding repository of reusable, standardized models (currently 60) for genomic, transcriptomic, and proteomic applications. Extensive validation on independent datasets demonstrates performance comparable to or exceeding that of state-of-the-art methods, including protein language models such as ESM-2. BioAutoMLFAST is available at https://bioautoml.icmc.usp.br/. This website is free and open to all users, and there is no login requirement.

## Introduction

The accurate functional and activity annotation of DNA, RNA, and protein sequences remains a significant challenge in the omics era, as high-throughput sequencing has produced data at a pace far exceeding experimental characterization capabilities (1, 2). Consequently, large fractions of biological sequences remain partially or entirely unannotated, limiting downstream biological interpretation and application (3). Computational approaches are therefore indispensable for translating sequence data into functional insight.

Traditionally, biological sequence analysis has relied on alignment-based methods that infer function through evolutionary conservation and homology. Tools such as BLAST (4) and profile-based approaches, including HMMER (5), underpin widely used resources such as Pfam (6) and KEGG (7). Despite their success, alignment-based methods depend on high-quality reference databases and may exhibit reduced sensitivity for short, highly divergent, or novel sequences lacking detectable homologs (8, 9).

Alignment-free, machine-learning-based approaches have emerged as powerful alternatives by learning predictive patterns directly from sequence-derived representations, enabling the analysis of divergent or previously uncharacterized sequences (10). These methods have been successfully applied across diverse tasks, including functional annotation, regulatory element detection, and phenotype prediction, with recent advances in deep learning and sequence language models further expanding representational capacity (11, 12). However, their broader adoption remains limited by the technical complexity of feature engineering, model selection, hyperparameter optimization, and rigorous evaluation (13, 14). Automated machine learning (AutoML) frameworks have been proposed to lower these barriers by automating key steps of the modeling pipeline (15–18). Despite their utility, existing AutoML tools for sequence analysis fail to address critical needs for robust and accessible biological discovery. First, some frameworks still lack support for raw biological sequences and require users to provide precomputed features. Although other platforms do accept sequences, the central limitation is their general inability to accommodate nucleotide and amino acid inputs simultaneously, together with limited emphasis on systematic exploration with feature engineering of sequence representations (e.g., *k*-mers, physicochemical properties, or embeddings), which is critical for capturing task-specific biological signals (16). In addition, most tools are ephemeral by design: models are typically trained for a single task and lack infrastructure to preserve them as standardized, reusable benchmarks. This absence of a persistent model repository severely impedes reproducible external validation and direct performance comparisons across studies (19).

Finally, the reliance on command-line interfaces creates a significant barrier for researchers without programming expertise (20). To address these limitations, we introduce BioAutoML-FAST, a **F**eature-based **A**utomated **S**ys**T**em, and publicly accessible, web-based AutoML platform for alignment-free biological sequence analysis. Table 1 compares our platform against representative frameworks, highlighting its unique integration of raw sequence processing, persistent benchmarking, and web-based accessibility.

**Table 1.** Comparison of representative automated machine-learning tools for biological sequence analysis. For each study, we report the availability of a publicly accessible dedicated web server, the presence of a model repository providing trained models for direct reuse, and support for automated feature engineering, defined as the systematic exploration of sequence representations. In addition, we indicate whether the tools support raw amino acid and nucleotide sequence inputs and classification and regression tasks. A check mark (✓) denotes explicit support for the corresponding capability. The symbol (∼) indicates limited support, e.g., web servers that report only performance metrics. Absence of a symbol indicates that the capability is not supported.

| Study | Web server | Model repository | Feature engineering | Amino acid | Nucleotide | Classification | Regression |
| --- | --- | --- | --- | --- | --- | --- | --- |
| AutoGenome (21) |  |  |  |  |  | ✓ | ✓ |
| iLearnPlus (15) | ✓ | ∼ |  | ✓ | ✓ | ✓ |  |
| JADBio (17) | ✓ | ✓ |  |  |  | ✓ | ✓ |
| BioAutoMATED (18) |  | ∼ |  | ✓ | ✓ | ✓ | ✓ |
| iM-Seeker (22) | ✓ |  |  |  | ✓ |  | ✓ |
| AutoPeptideML (23) | ∼ | ✓ |  | ✓ |  | ✓ |  |
| AutoProteinEngine (24) |  |  | ∼ | ✓ |  | ✓ | ✓ |
| aMLProt (25) |  |  |  | ✓ |  | ✓ | ✓ |
| BioAutoML (16) |  |  | ✓ | ✓ | ✓ | ✓ |  |
| BioAutoML-FAST | ✓ | ✓ | ✓ | ✓ | ✓ | ✓ | ✓ |

The platform not only builds on our earlier BioAutoML framework (16) but also substantially extends this foundation by operationalizing persistent benchmarking as a first-class design principle. By deploying the resource as a production-ready web resource within a broader, methodologically integrated infrastructure, we complete the final step of Chou’s five-step rule (26, 27). Our platform integrates automated feature extraction, engineering, model training, evaluation, and visualization within a unified interface. It supports end-to-end analysis of both nucleotide and amino acid sequences for classification and regression tasks, with optional data encryption. Beyond automated analysis, BioAutoML-FAST serves as a community benchmarking resource by preserving trained models as persistent, reusable benchmarks evaluated under standardized protocols in a curated, expandable repository. By combining accessible analysis with standardized, persistent benchmarking, BioAutoML-FAST provides a reproducible and scalable computational infrastructure for the life sciences.

## Materials and Methods

This section outlines the sequence representations, automated modeling strategy, benchmark datasets, and evaluation protocols used in BioAutoML-FAST. It summarizes the numerical descriptors for nucleotide and amino acid encoding; the fully automated pipeline for selecting optimal descriptor–algorithm combinations; performance metrics for classification and regression; and the curation of prior datasets supporting model training, external validation, and the creation of a reusable repository.

### Biological sequence representations

BioAutoMLFAST systematically explores a comprehensive suite of sequence feature representations (descriptors) to identify the most informative ones for a given dataset. The platform incorporates 20 distinct descriptors for nucleotide sequences and 23 for amino acid sequences. These descriptors are derived from established feature extraction packages in the literature, including Chen et al. (15), Liu et al. (28), Müller et al. (29), Bonidia et al. (30). Table 2 describes the descriptors used by the platform for nucleotide sequences, while Table 3 describes the descriptors for amino acid sequences. The selection of the feature representations was guided by three criteria: frequent use in prior literature, manageable feature dimensionality, and reasonable computational cost.

**Table 2.** Descriptors for numerical representation of nucleotide sequences. For each descriptor, the short name, full description, feature dimension (total number of extracted numerical features), and primary reference are provided.

| Short name | Descriptor | Feature dimension | Reference |
| --- | --- | --- | --- |
| NAC | Nucleic acid composition (NAC) | 4 | (30) |
| DNC | Dinucleotide composition (DNC) | 16 | (30) |
| TNC | Trinucleotide composition (TNC) | 64 | (30) |
| kGap_di | Xmer k-Spaced Ymer composition (kGap) – frequency of 2 after 1-gap | 64 | (30) |
| kGap_tri | Xmer k-Spaced Ymer composition (kGap) – frequency of 3 after 1-gap | 256 | (30) |
| ORF | Open reading frame-based features | 10 | (30) |
| Fickett | Fickett score based on positional nucleotide features | 2 | (30) |
| Shannon | Shannon’s entropy from 1-mer to 5-mer | 5 | (30) |
| Tsallis | Tsallis’ entropy from 1-mer to 5-mer with $q = 2.3$ | 5 | (30) |
| FourierBinary | Binary numerical mapping using Fourier transform | 19 | (30) |
| FourierComplex | Complex numerical mapping using Fourier transform | 19 | (30) |
| Revkmer | Reverse compliment kmer | 44 | (28) |
| PseDNC | Combining dinucleotide composition and global sequence-order effects | 19 | (28) |
| PseKNC | Improving PseDNC by incorporating k-tuple nucleotide composition | 65 | (28) |
| SC-PseDNC | Combining dinucleotide composition and global sequence-order effects by series correlation | 54 | (28) |
| SC-PseTNC | Combining trinucleotide composition and global sequence-order effects by series correlation | 88 | (28) |
| DAC | Incorporating the correlation of the same property between two dinucleotides | 76 | (28) |
| TAC | Incorporating the correlation of the same property between two trinucleotides | 24 | (28) |
| TCC | Incorporating the correlation of the different properties between two trinucleotides | 264 | (28) |
| TACC | Combination of TAC and TCC | 288 | (28) |

**Table 3.** Descriptors for numerical representation of amino acid sequences. For each descriptor, the short name, full description, feature dimension (total number of extracted numerical features), and primary reference are provided.

| Short name | Descriptor | Feature dimension | Reference |
| --- | --- | --- | --- |
| AAC | Amino acid composition (AAC) | 20 | (30) |
| DPC | Dipeptides composition (DPC) | 400 | (30) |
| kGap | Xmer k-Spaced Ymer composition (kGap) – frequency of 1 after 1-gap | 400 | (30) |
| CKSAAP | Composition of k-spaced amino acid pairs | 2,400 | (15) |
| DDE | Dipeptide deviation from expected mean | 400 | (15) |
| GAAC | Grouped amino acid composition | 5 | (15) |
| CKSAAGP | Composition of k-spaced amino acid group pairs | 150 | (15) |
| GDPC | Grouped dipeptide composition | 25 | (15) |
| GTPC | Grouped tripeptide composition | 125 | (15) |
| CTDC | Composition (CTDC) | 39 | (15) |
| CTDT | Transition (CTDT) | 39 | (15) |
| CTDD | Distribution (CTDD) | 195 | (15) |
| CTriad | Conjoint triad | 343 | (15) |
| KSCTriad | Conjoint k-spaced triad | 343 | (15) |
| Global | Global one-dimensional peptide descriptors | 10 | (29) |
| Peptide | AA scale based global or convoluted descriptors (auto-/cross-correlated) | 16 | (29) |
| Fourier_Integer | Integer numerical mapping using Fourier transform | 19 | (30) |
| Fourier_EIIP | Electron-ion interaction potential numerical mapping using Fourier transform | 19 | (30) |
| Shannon | Shannon’s entropy from 1-mer to 5-mer | 5 | (30) |
| Tsallis_23 | Tsallis’s entropy from 1-mer to 5-mer with $q = 2.3$ | 5 | (30) |
| Tsallis_30 | Tsallis’s entropy from 1-mer to 5-mer with $q = 3.0$ | 5 | (30) |
| Tsallis_40 | Tsallis’s entropy from 1-mer to 5-mer with $q = 4.0$ | 5 | (30) |
| ComplexNetworks | Complex network features from 1-mer to 5-mer | 78 | (30) |

### Automated machine learning

BioAutoML-FAST implements a full AutoML pipeline that minimizes user intervention and builds upon the original BioAutoML framework (16). Unlike platforms such as iLearnPlus (15) which require manual selection of parameters or descriptors, BioAutoML-FAST automates the entire process from feature engineering to algorithm selection. To navigate the exponentially large combinatorial search space efficiently, the system employs the Optuna framework with the Tree-structured Parzen Estimator (TPE) (31). This end-to-end automation ensures users can obtain high-performing models without requiring expertise in parameter tuning or descriptor selection.

The pipeline employs a two-step Bayesian optimization strategy (Figure 4). In the first step, the framework identifies optimal feature descriptors using a fixed machine-learning algorithm (e.g., LightGBM) across 200 optimization trials, allocating 80 trials for early stopping if model performance fails to improve by at least 0.001. In the second step, the selected descriptors are used to jointly optimize the learning algorithm and its hyperparameters across 150 trials. Both steps aim to maximize the Matthews Correlation Coefficient (MCC) for classification or minimize Root Mean Squared Error (RMSE) for regression, using 5-fold cross-validation. The number of trials was determined by the size of the search space and the trade-off between predictive performance and computational cost.

Finally, the resulting optimal model is subjected to a more rigorous evaluation, including a refined 10-fold cross-validation to obtain a robust performance estimate and, when available, validation on an independent external test set.

### Performance evaluation

Model performance was assessed using metrics routinely adopted in supervised biological sequence prediction and general machine-learning benchmarks (32, 33). For binary classification, we report Sensitivity (Sn), Specificity (Sp), Accuracy (Acc), MCC, and the Area Under the Receiver Operating Characteristic Curve (AUC), which together capture true class assignment and threshold-independent discriminative ability. For multiclass classification, we report Sn, Sp, Acc, and MCC. Sensitivity and specificity were computed in a one-versus-rest scheme and summarized using macro-averaging, the standard strategy in bioinformatics for providing balanced assessment when class distributions are uneven. For regression, we report Mean Absolute Error (MAE), Mean Squared Error (MSE), RMSE, and the Coefficient of Determination (*R*^2^), the standard measures for continuous prediction performance that capture absolute and squared deviations and explained variance.

### Benchmark datasets

The benchmark datasets used in BioAutoML-FAST were collected from previously published studies covering a broad spectrum of biological sequence analysis tasks, including genomic, transcriptomic, and proteomic applications. In total, the platform incorporates 60 benchmark datasets, comprising 50 binary classification, seven multiclass classification, and three regression problems, as summarized in Table 4. These datasets include both nucleotide and amino acid sequences, with 24 nucleotide-based datasets and 36 amino acid–based datasets. For each benchmark, the original data splits defined in the source studies were preserved whenever available, with training and external test datasets clearly distinguished. In cases where external test sets were not reported in the original publications, models were trained exclusively on the available training data, and the absence of independent test data is explicitly indicated. This strategy ensures faithful reproduction of the experimental conditions under which the datasets were initially introduced, while maintaining transparency regarding dataset availability.

**Table 4.** Benchmark datasets used to train and evaluate the models available in BioAutoML-FAST. The table summarizes the prediction task (binary classification, multiclass classification, or regression), sequence type, and the size of the training and external test datasets for each model, along with the original reference from which the data were derived. A dash (–) indicates that the dataset was not available or not reported in the original study.

| Model | Task | Sequence type | Training dataset | Test dataset | Reference |
| --- | --- | --- | --- | --- | --- |
| Model 1: antibody peptides | Binary | Amino acid | 8,959 positives / 54,441 negatives | - | (48) |
| Model 2: anticancer peptides | Binary | Amino acid | 250 positives / 250 negatives | 82 positives / 82 negatives | (49) |
| Model 3: anticancer peptides | Binary | Amino acid | 138 positives / 206 negatives | - | (50) |
| Model 4: anticancer peptides | Binary | Amino acid | 776 positives / 776 negatives | 194 positives / 194 negatives | (51) |
| Model 5: anticancer peptides | Binary | Amino acid | 689 positives / 689 negatives | 172 positives / 172 negatives | (51) |
| Model 6: anti-coronavirus | Binary | Amino acid | 109 positives / 356 negatives | - | (52) |
| Model 7: anti-coronavirus and random non-secretory proteins | Binary | Amino acid | 109 positives / 539 negatives | - | (52) |
| Model 8: antifungal peptides | Binary | Amino acid | 778 positives / 778 negatives | 215 positives / 215 negatives | (53) |
| Model 9: anti-hypertensive peptides | Binary | Amino acid | 913 positives / 913 negatives | 386 positives / 386 negatives | (54) |
| Model 10: antimalarial peptides | Binary | Amino acid | 111 positives / 542 negatives | 28 positives / 135 negatives | (55) |
| Model 11: antimalarial peptides | Binary | Amino acid | 111 positives / 1,708 negatives | 28 positives / 427 negatives | (55) |
| Model 12: antimicrobial peptides | Binary | Amino acid | 1,686 positives / 16,428 negatives | 723 positives / 7,041 negatives | (56) |
| Model 13: antimicrobial peptides | Binary | Amino acid | 879 positives / 2,405 negatives | 920 positives / 920 negatives | (57) |
| Model 14: antimicrobial peptides | Binary | Amino acid | 3,876 positives / 9,552 negatives | 2,584 positives / 6,369 negatives | (58) |
| Model 15: anti-MRSA strains peptides | Binary | Amino acid | 118 positives / 678 negatives | 30 positives / 169 negatives | (59) |
| Model 16: antioxidant proteins | Binary | Amino acid | 249 positives / 1,531 negatives | 74 positives / 392 negatives | (60, 61) |
| Model 17: anti-parasitic peptides | Binary | Amino acid | 255 positives / 1,863 negatives | 46 positives / 46 negatives | (62) |
| Model 18: antiviral | Binary | Amino acid | 520 positives / 379 negatives | 57 positives / 41 negatives | (52) |

Table 4. (continued)
| Model | Task | Sequence type | Training dataset | Test dataset | Reference |
| --- | --- | --- | --- | --- | --- |
| Model 19: antiviral and random non-secretory proteins | Binary | Amino acid | 520 positives / 539 negatives | 57 positives / 58 negatives | (52) |
| Model 20: antiviral peptides | Binary | Amino acid | 2,321 positives / 2,321 negatives | 623 positives / 623 negatives | (53) |
| Model 21: bitter peptides | Binary | Amino acid | 256 positives / 256 negatives | 64 positives / 64 negatives | (63) |
| Model 22: blood-brain barrier peptides | Binary | Amino acid | 100 positives / 100 negatives | 19 positives / 19 negatives | (64) |
| Model 23: DNA-binding proteins | Binary | Amino acid | 524 positives / 550 negatives | 93 positives / 90 negatives | (65, 66) |
| Model 24: DNA-binding proteins | Binary | Amino acid | 146 positives / 250 negatives | - | (67) |
| Model 25: DNase I hypersensitive sites | Binary | Nucleotide | 280 positives / 737 negatives | - | (68) |
| Model 26: DPP IV inhibitory peptides | Binary | Amino acid | 532 positives / 532 negatives | 133 positives / 133 negatives | (69) |
| Model 27: hemolytic peptides | Binary | Amino acid | 442 positives / 442 negatives | 110 positives / 110 negatives | (70) |
| Model 28: recombination spots | Binary | Nucleotide | 478 positives / 572 negatives | - | (71, 72) |
| Model 29: lncRNA (*Homo sapiens*) | Binary | Nucleotide | 8,000 positives / 8,000 negatives | 2,500 positives / 2,500 negatives | (73) |
| Model 30: lncRNA (*Mus musculus*) | Binary | Nucleotide | 4,200 positives / 4,200 negatives | 1,800 positives / 1,800 negatives | (73) |
| Model 31: lncRNA (*Triticum aestivum*) | Binary | Nucleotide | 4,000 positives / 4,000 negatives | 2,000 positives / 2,000 negatives | (73) |
| Model 32: lncRNA (*Zea mays*) | Binary | Nucleotide | 18,000 positives / 18,000 negatives | - | (74) |
| Model 33: m5C sites (*Arabidopsis thaliana*) | Binary | Nucleotide | 5,289 positives / 5,289 negatives | 1,000 positives / 1,000 negatives | (75) |
| Model 34: m5C sites (*Homo sapiens*) | Binary | Nucleotide | 120 positives / 120 negatives | - | (76) |
| Model 35: m5C sites (*Mus musculus*) | Binary | Nucleotide | 97 positives / 97 negatives | - | (75) |
| Model 36: m5C sites (*Saccharomyces cerevisiae*) | Binary | Nucleotide | 211 positives / 211 negatives | - | (75) |
| Model 37: neuropeptides | Binary | Amino acid | 1,940 positives / 1,940 negatives | 485 positives / 485 negatives | (77, 78) |

Table 4. (continued)
| Model | Task | Sequence type | Training dataset | Test dataset | Reference |
| --- | --- | --- | --- | --- | --- |
| Model 38: non-classical secreted proteins | Binary | Amino acid | 141 positives / 446 negatives | 34 positives / 34 negatives | (79) |
| Model 39: peptide toxicity | Binary | Amino acid | 1,663 positives / 1,621 negatives | 269 positives / 311 negatives | (80) |
| Model 40: phage virion proteins | Binary | Amino acid | 250 positives / 250 negatives | 63 positives / 63 negatives | (81) |
| Model 41: proinflammatory peptides | Binary | Amino acid | 607 positives / 1,098 negatives | 134 positives / 156 negatives | (82) |
| Model 42: protein lysine crotonylation sites | Binary | Amino acid | 2,742 positives / 29,676 negatives | 711 positives / 7,458 negatives | (83) |
| Model 43: quorum-sensing peptides | Binary | Amino acid | 200 positives / 200 negatives | 20 positives / 20 negatives | (84) |
| Model 44: real microRNA precursors | Binary | Nucleotide | 1,612 positives / 1,612 negatives | - | (85) |
| Model 45: ribosome binding site sequences | Binary | Nucleotide | 25,000 positives / 25,000 negatives | - | (86) |
| Model 46: sigma70 promoters | Binary | Nucleotide | 741 positives / 1,400 negatives | - | (87) |
| Model 47: small non-coding RNA and shuffled sequences | Binary | Nucleotide | 182 positives / 182 negatives | - | (88) |
| Model 48: toehold switch sequences | Binary | Nucleotide | 45,758 positives / 45,776 negatives | - | (89) |
| Model 49: Tumor T cell antigens | Binary | Amino acid | 470 positives / 318 negatives | 122 positives / 75 negatives | (90) |
| Model 50: umami peptides | Binary | Amino acid | 112 positives / 241 negatives | 28 positives / 61 negatives | (91) |
| Model 51: lncRNAs subcellular localization - training set with 4 classes | Multiclass | Nucleotide | Cytoplasm: 292 / Cytosol: 91 / Exosome: 25 / Nucleus: 149 / Ribosome: 43 | Cytoplasm: 199 / Exosome: 16 / Nucleus: 82 / Ribosome: 99 | (92) |
| Model 52: lncRNAs subcellular localization - training set with 5 classes | Multiclass | Nucleotide | Cytoplasm: 417 / Exosome: 30 / Nucleus: 153 / Ribosome: 43 | Cytoplasm: 199 / Exosome: 16 / Nucleus: 82 / Ribosome: 99 | (92) |

Table 4. (continued)
| Model | Task | Sequence type | Training dataset | Test dataset | Reference |
| --- | --- | --- | --- | --- | --- |
| Model 53: mRNA subcellular localization | Multiclass | Nucleotide | Cytoplasm: 5,310 / Endoplasmic reticulum: 1,185 / Extracellular region: 710 / Mitochondria: 350 / Nucleus: 4,855 | Cytoplasm: 1,066 / Endoplasmic reticulum: 241 / Extracellular region: 145 / Mitochondria: 71 / Nucleus: 976 | (93) |
| Model 54: non-coding RNA - 4 classes | Multiclass | Nucleotide | Cis-reg: 246 / rRNA: 242 / sRNA: 497 / rRNA: 581 | Cis-reg: 169 / rRNA: 325 / sRNA: 119 / rRNA: 50 | (94) |
| Model 55: non-coding RNA - 8 classes | Multiclass | Nucleotide | miRNA: 212 / mRNA: 514 / pre-miRNA: 203 / rRNA: 593 / snoRNA: 143 / snRNA: 91 / tmRNA: 282 / tRNA: 358 | miRNA: 52 / mRNA: 188 / pre-miRNA: 50 / rRNA: 148 / snoRNA: 35 / snRNA: 22 / tmRNA: 70 / tRNA: 89 | (16) |
| Model 56: non-coding RNA - E. coli K12 | Multiclass | Nucleotide | rRNA: 32 / sRNA: 133 / tRNA: 40 | rRNA: 8 / sRNA: 33 / tRNA: 10 | (16) |
| Model 57: non-coding RNA - Multiple bacterial phyla | Multiclass | Nucleotide | rRNA: 144 / sRNA: 309 / tRNA: 329 | rRNA: 103 / sRNA: 179 / tRNA: 267 | (16) |
| Model 58: antibody peptides | Regression | Amino acid | peptides: 67,769 | - | (48) |
| Model 59: ribosome binding site sequences | Regression | Nucleotide | rbs: 50,000 | - | (86) |
| Model 60: toehold switch sequences | Regression | Nucleotide | toeholds: 91,534 | - | (89) |

### Server implementation

BioAutoML-FAST is developed in Python (34) and adopts Streamlit (35) as the web application framework. Input and output files are handled through Biopython (36), while data parsing and manipulation rely on Pandas (37). Visualizations are produced using Matplotlib (38) and Plotly (39), with Seaborn (40) employed for additional graphical representations. The machine-learning models are built upon scikit-learn (41), XGBoost (42), and Light-GBM (43). To support interpretable predictions, the platform integrates the SHAP library (44) in combination with custom Python scripts. Task scheduling and asynchronous execution are coordinated by Redis (45), which manages the queue system. Detailed information on software versions and dependencies is available in the code repository, as indicated in the Data Availability section.

## Results

This section provides an overview of the server’s capabilities and presents extensive comparisons across benchmark datasets, illustrating its functionality and demonstrating its general applicability.

### Web server usage

BioAutoML-FAST provides four main interfaces (Figure 1), starting with the Home interface. Here, users can either train a new model from scratch or load a trained model by providing FASTA files for sequence data or a CSV file for structured data. The user specifies the task type (classification or regression), data type (amino acid sequences, nucleotide sequences, or structured data), and the evaluation method: (i) training without a test set, (ii) with a labeled test set, or (iii) with an unlabeled prediction set. Optional hyperparameter tuning is available for improved robustness, though it increases training time. Users can optionally provide an email address to be notified on job completion and encrypt their submission with a password to ensure secure access. The submission generates an asynchronous job-based analysis.

**Fig. 1.**
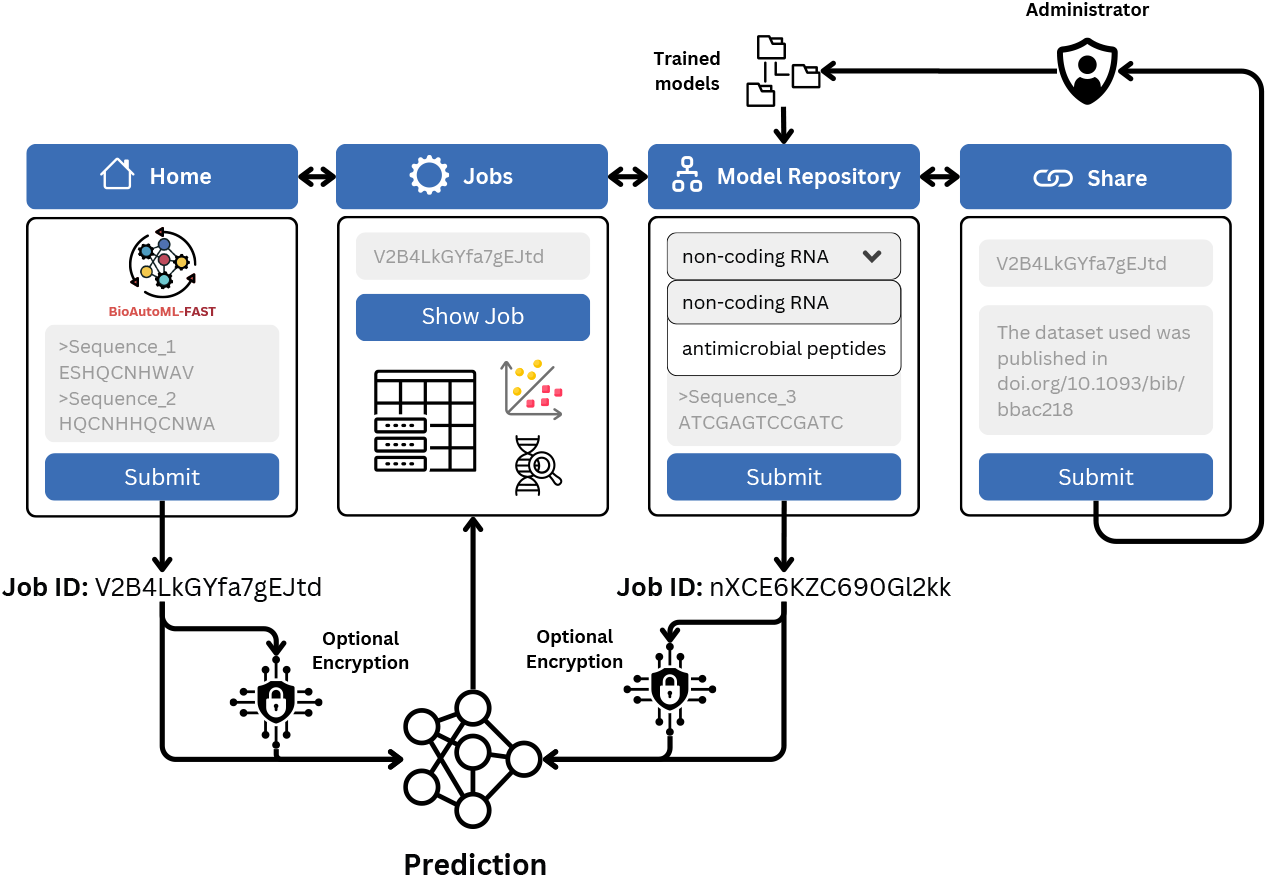
Overview of the BioAutoML-FAST platform. The platform provides four main functionalities: (1) Home – submit nucleotide sequences, amino acid sequences or structured data to train a model from scratch or load a model; (2) Jobs – view submission results with multiple interactive visualizations; (3) Model Repository – access to trained models for diverse tasks; (4) Share – share a trained model, requesting its addition to the repository.

Users access submission results via the Jobs interface by entering their Job ID and, if applicable, a decryption password. This interface serves as a comprehensive dashboard for multifaceted analysis. It allows users to download trained models and metadata, review performance metrics for training and test datasets, and examine class-specific probability predictions. Furthermore, the interface provides insights into feature importance using tree-based methods or SHAP values, feature distributions, and Pearson or Spearman correlations. Finally, it offers 3D dataset visualization using dimensionality reduction techniques such as PCA, t-SNE (46), and UMAP (47).

In the Model Repository interface, users can select from 60 models trained for various tasks, such as anticancer peptide prediction and non-coding RNA classification. For each model, a preview is available for the predictive tasks, data type, possible prediction labels, dataset composition, and the reference paper from which the model’s dataset originates. Users can submit unlabeled data for prediction and, optionally, provide their email address for notifications and a password for encryption. This submission generates an asynchronous, job-based analysis similar to the Home interface. The Share interface allows users to submit their trained model to the platform for inclusion in the Model Repository, subject to administrative review. To do so, users provide the associated Job ID, an optional email address, and a reference to the source of the training dataset.

### Performance Evaluation on Benchmark Datasets

To rigorously assess BioAutoML-FAST, we evaluated models on 60 diverse benchmark datasets (Table 4) spanning nucleotide and amino acid tasks, including binary and multiclass classification and regression. For each dataset, the automated pipeline identified optimal feature-algorithm combinations, with experiments repeated five times to quantify stability. We compared the resulting models against state-of-the-art predictors using 152 training and 179 testing metrics, ensuring a robust evaluation of predictive performance and generalizability.

Performance metrics for BioAutoML-FAST are reported as the mean *±* standard deviation across 5 repeated runs to quantify performance variability. For comparative analysis, we assessed whether the single-value results reported by SOTA studies fell within the performance intervals of our models. This interval-based assessment was adopted instead of formal statistical hypothesis testing (e.g., paired t-tests) because the raw prediction data and cross-validation variance for many external baselines were not publicly available, precluding direct statistical comparison. Under this criterion, BioAutoML-FAST matched or outperformed reported SOTA predictors on more than half of the training metrics (90/152) and nearly half of the external validation metrics (87/179). Notably, BioAutoML-FAST repeatedly outperformed studies using protein language models (pLMs) such as ESM-2, as shown in Models 10, 11, 14, 15, 20, 26, and 49. Among the remaining cases, performance was highly competitive: 102 training metrics and 135 testing metrics were within 5% of the corresponding SOTA results. Detailed comparative benchmarks are provided for binary classification (Table 5), multiclass classification (Table 6), and regression tasks (Table 7).

**Table 5.**
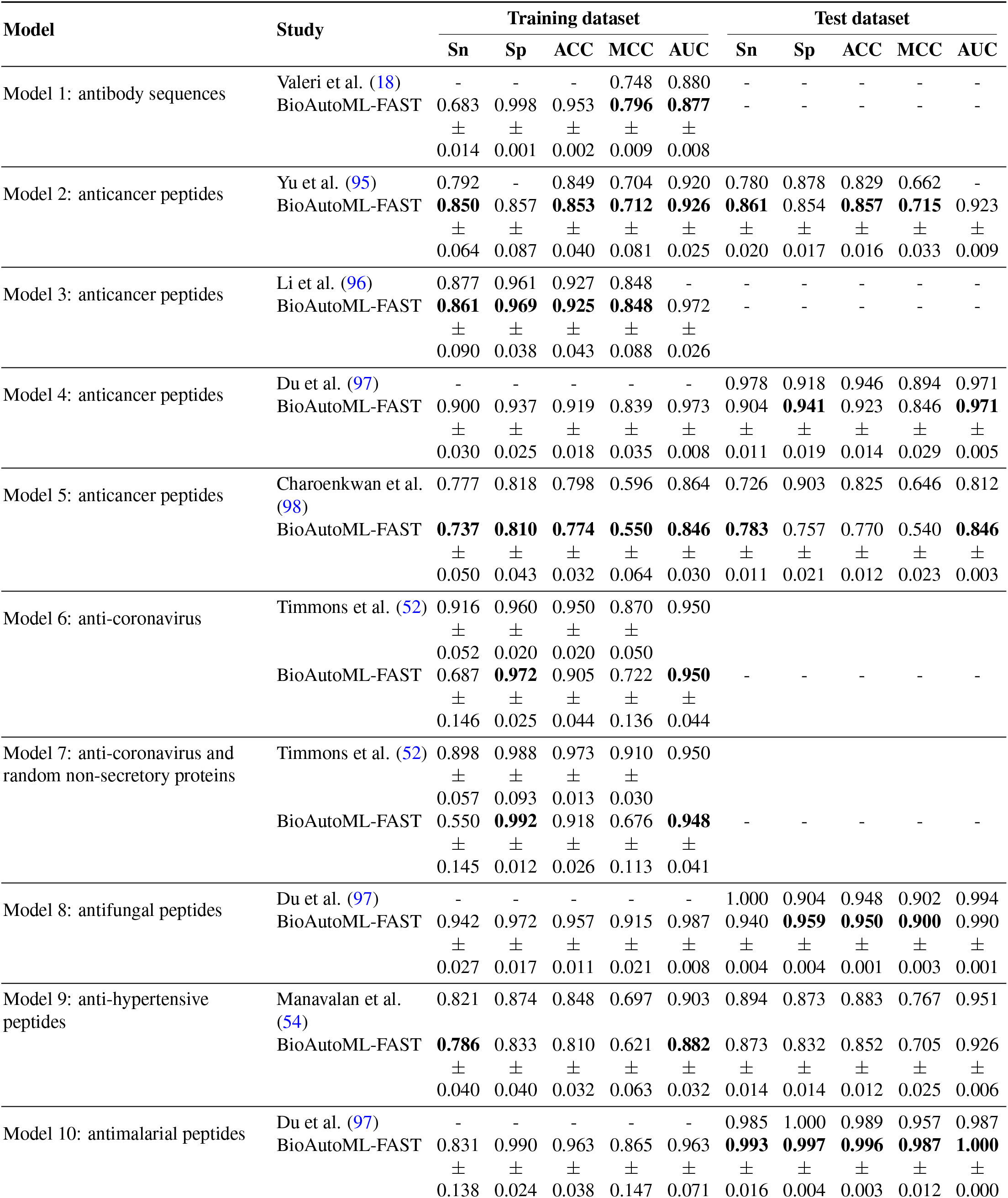
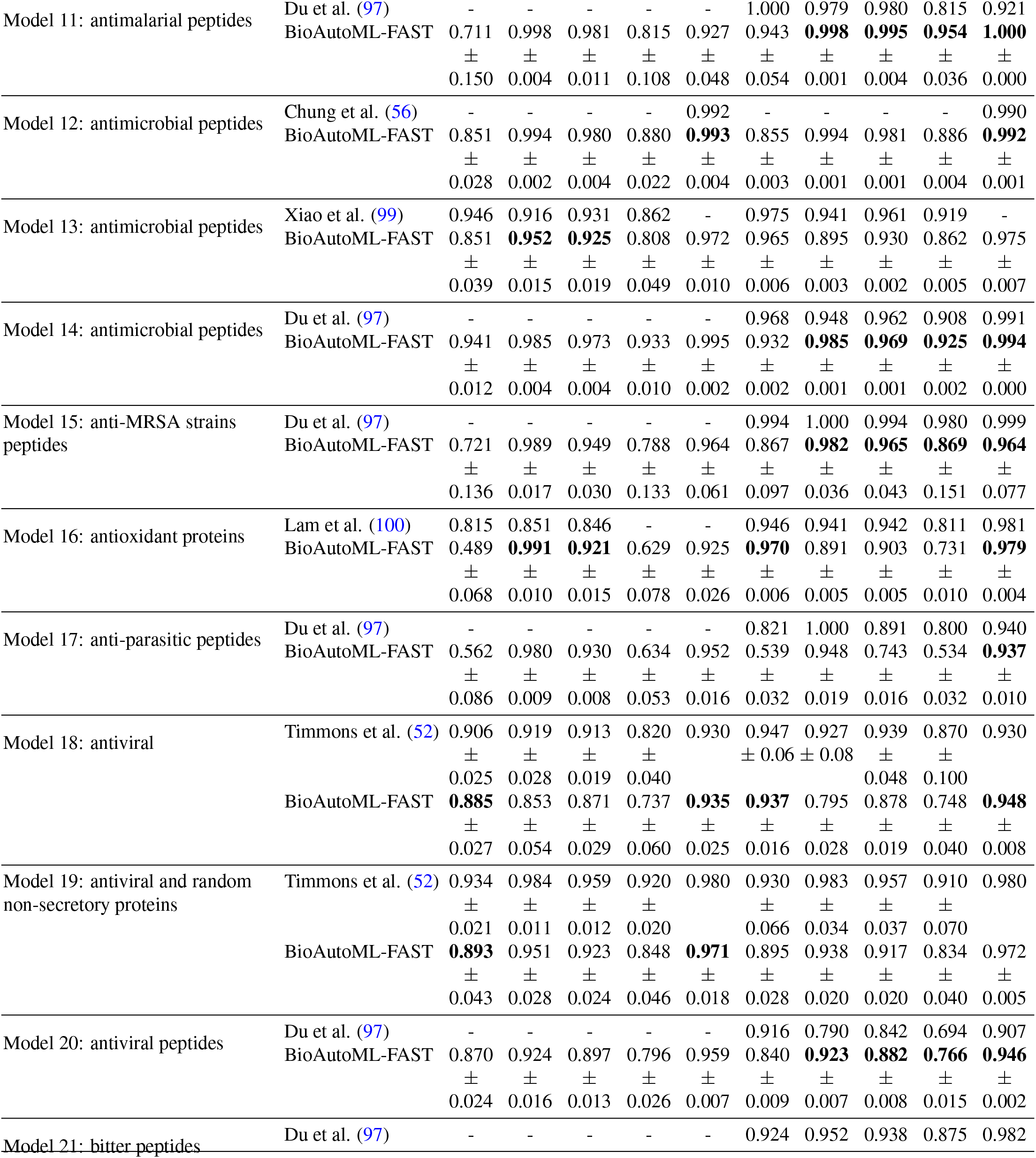
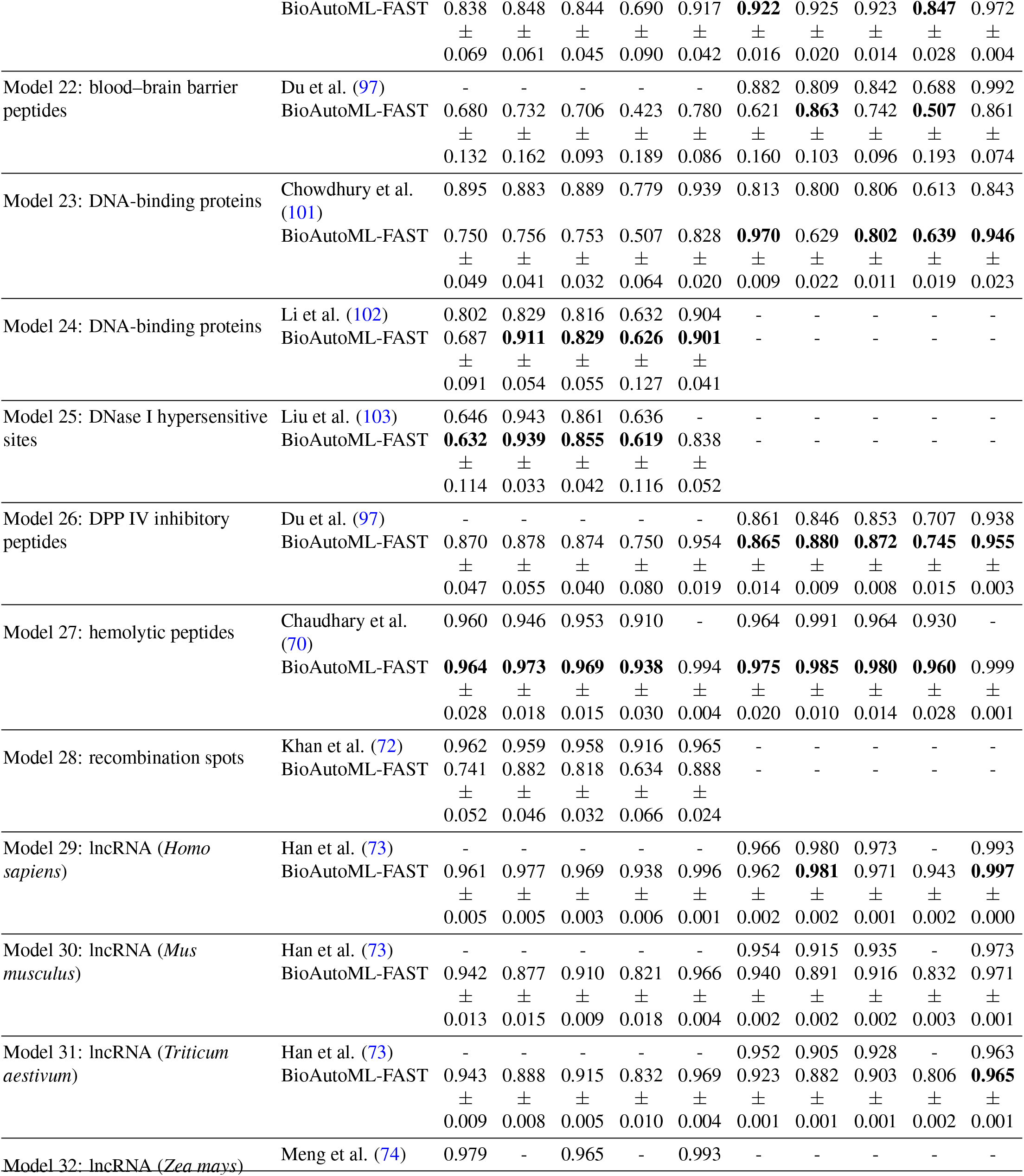
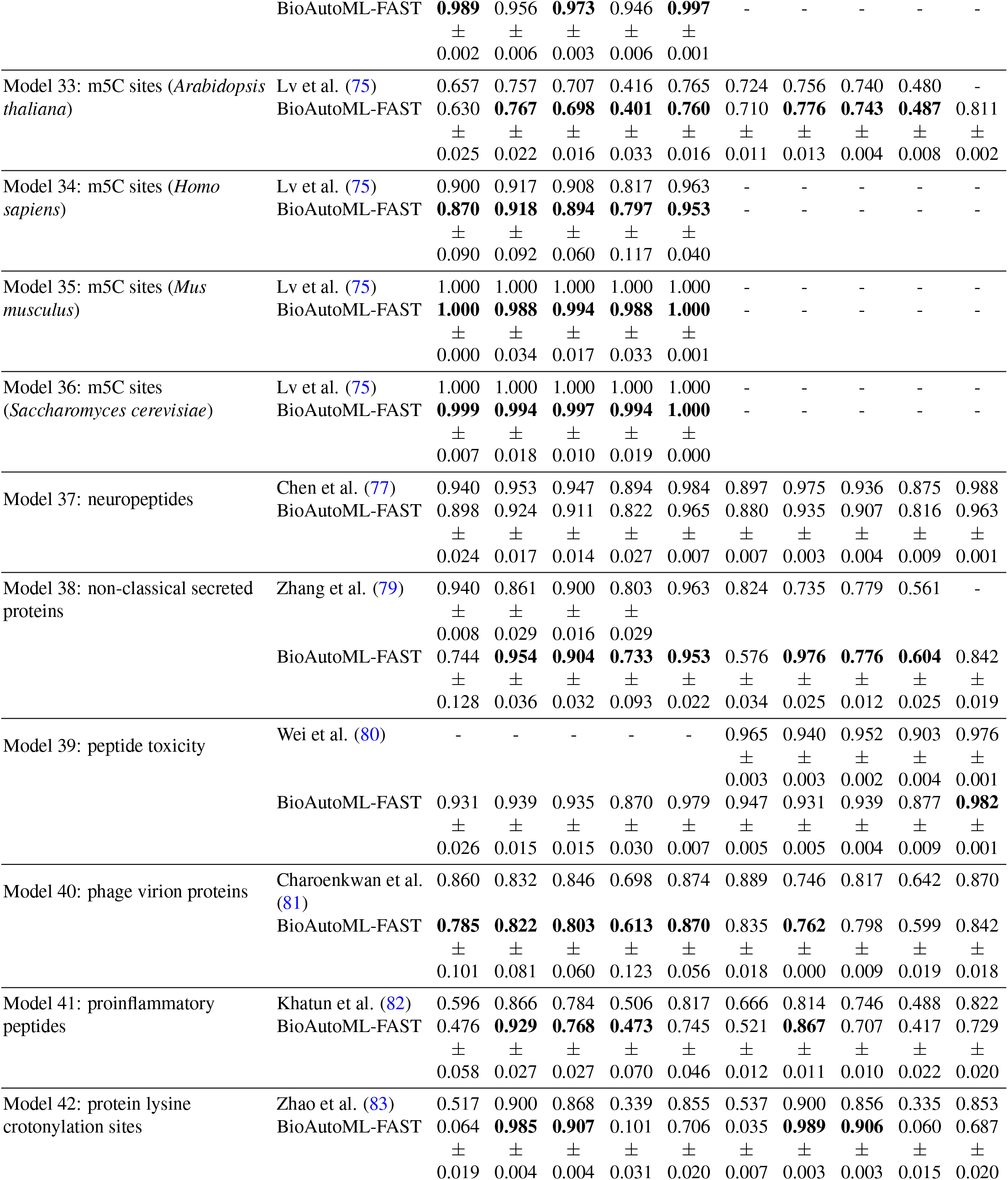
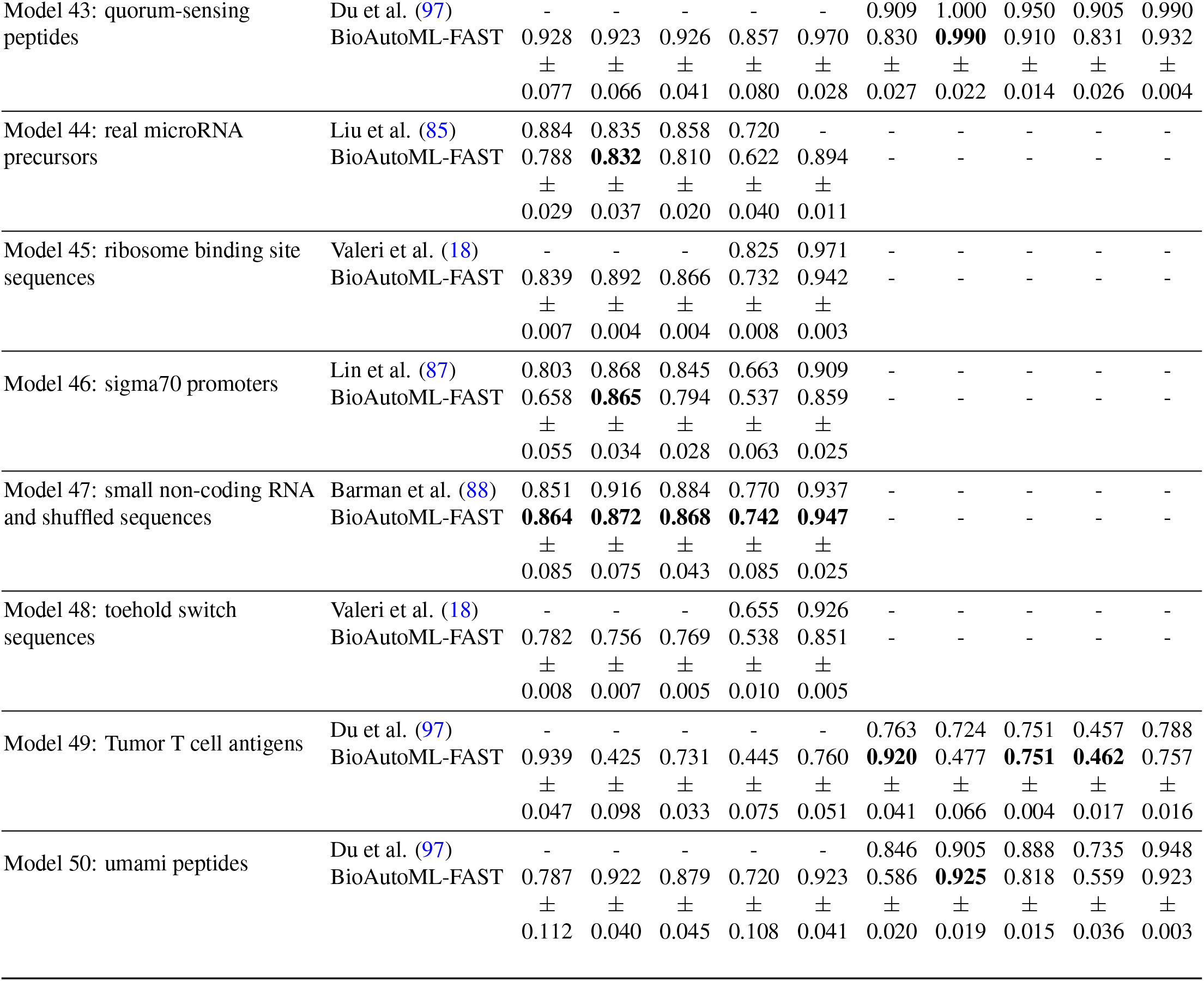
Performance comparison of binary models across datasets. Metrics include Sensitivity (Sn), Specificity (Sp), Accuracy (ACC), Matthews Correlation Coefficient (MCC), and Area Under the Curve (AUC). Dashes (-) indicate metrics that were not reported in the baseline study. Metrics for which BioAutoML-FAST exhibits comparable performance (within one standard deviation of the best reported value) or superior results are highlighted in bold.

**Table 6.** Performance comparison of multiclass models across datasets. Metrics include Sensitivity (Sn), Specificity (Sp), Accuracy (ACC), Matthews Correlation Coefficient (MCC), and Area Under the Curve (AUC). Dashes (-) indicate metrics that were not reported in the baseline study. Metrics for which BioAutoML-FAST exhibits comparable performance (within one standard deviation of the best reported value) or superior results are highlighted in bold.

| Model | Study | Training dataset |  |  |  | Test dataset |  |  |  |
| --- | --- | --- | --- | --- | --- | --- | --- | --- | --- |
|  |  | Sn | Sp | ACC | MCC | Sn | Sp | ACC | MCC |
| Model 51: lncRNAs subcellular localization - training set with 4 classes | Cai et al. (92) | 0.945 | 0.982 | 0.942 | 0.928 | 0.463 | - | 0.462 | - |
|  | BioAutoML-FAST | 0.332 | 0.787 | 0.663 | 0.209 | 0.298 | 0.765 | **0.459** | 0.069 |
|  |  | ± | ± | ± | ± | ± | ± | ± | ± |
|  |  | 0.077 | 0.026 | 0.040 | 0.116 | 0.024 | 0.011 | 0.033 | 0.051 |
| Model 52: lncRNAs subcellular localization - training set with 5 classes | Cai et al. (92) | 0.933 | - | 0.934 | - | 0.463 | - | 0.462 | - |
|  | BioAutoML-FAST | 0.330 | 0.861 | 0.574 | 0.321 | 0.418 | 0.789 | **0.482** | 0.182 |
|  |  | ± | ± | ± | ± | ± | ± | ± | ± |
|  |  | 0.050 | 0.019 | 0.059 | 0.102 | 0.016 | 0.006 | 0.018 | 0.025 |
| Model 53: mRNA subcellular localization | Musleh et al. (93) | 0.840 | 0.829 | 0.834 | - | 0.792 | 0.759 | 0.748 | - |
|  | BioAutoML-FAST | 0.707 | **0.940** | 0.821 | 0.718 | 0.447 | **0.820** | 0.272 | 0.128 |
|  |  | ± | ± | ± | ± | ± | ± | ± | ± |
|  |  | 0.021 | 0.004 | 0.012 | 0.019 | 0.012 | 0.002 | 0.011 | 0.008 |
| Model 54: non-coding RNA - 4 classes | Avila et al. (94) | - | - | - | - | 0.921 | - | 0.928 | - |
|  | BioAutoML-FAST | 0.910 | 0.976 | 0.929 | 0.900 | 0.896 | 0.971 | 0.908 | 0.862 |
|  |  | ± | ± | ± | ± | ± | ± | ± | ± |
|  |  | 0.022 | 0.006 | 0.016 | 0.022 | 0.009 | 0.003 | 0.008 | 0.012 |
| Model 55: non-coding RNA - 8 classes | Bonidia et al. (16) | - | - | - | - | 0.776 | 0.976 | 0.855 | 0.822 |
|  | BioAutoML-FAST | 0.801 | 0.983 | 0.877 | 0.854 | **0.799** | **0.985** | **0.894** | **0.870** |
|  |  | ± | ± | ± | ± | ± | ± | ± | ± |
|  |  | 0.029 | 0.002 | 0.017 | 0.021 | 0.009 | 0.000 | 0.003 | 0.003 |
| Model 56: non-coding RNA - E. coli K12 | Bonidia et al. (16) | - | - | - | - | 0.990 | 0.992 | 0.980 | 0.964 |
|  | BioAutoML-FAST | 0.972 | 0.983 | 0.979 | 0.961 | **0.987** | **0.993** | **0.992** | **0.985** |
|  |  | ± | ± | ± | ± | ± | ± | ± | ± |
|  |  | 0.062 | 0.036 | 0.039 | 0.073 | 0.018 | 0.010 | 0.011 | 0.021 |
| Model 57: non-coding RNA - Multiple bacterial phyla | Bonidia et al. (16) | - | - | - | - | 0.979 | 0.990 | 0.982 | 0.971 |
|  | BioAutoML-FAST | 0.983 | 0.991 | 0.983 | 0.974 | 0.974 | **0.989** | **0.979** | **0.967** |
|  |  | ± | ± | ± | ± | ± | ± | ± | ± |
|  |  | 0.031 | 0.018 | 0.034 | 0.051 | 0.003 | 0.002 | 0.004 | 0.006 |

**Table 7.** Performance comparison of regression models across datasets. Metrics include Mean Absolute Error (MAE), Mean Squared Error (MSE), Root Mean Squared Error (RMSE), and the coefficient of determination (*R*^2^). Dashes (-) indicate metrics that were not reported in the baseline study. Metrics for which BioAutoML-FAST exhibits comparable performance (within one standard deviation of the best reported value) or superior results are highlighted in bold.

| Model | Study | Training dataset |  |  |  | Test dataset |  |  |  |
| --- | --- | --- | --- | --- | --- | --- | --- | --- | --- |
|  |  | MAE | MSE | RMSE | R2 | MAE | MSE | RMSE | R2 |
| Model 58 - antibody peptides | Valeri et al. (18) | 0.382 | - | - | 0.452 | - | - | - | - |
|  | BioAutoML-FAST | **0.382** | 0.233 | 0.483 | **0.448** | - | - | - | - |
| | | ± | ± | ± | ± | | | | |
|  |  | 0.003 | 0.004 | 0.004 | 0.009 |  |  |  |  |
| Model 59 - ribosome binding site sequences | Valeri et al. (18) | 0.067 | - | - | 0.854 | - | - | - | - |
|  | BioAutoML-FAST | 0.076 | 0.012 | 0.111 | **0.856** | - | - | - | - |
| | | ± | ± | ± | ± | | | | |
|  |  | 0.002 | 0.001 | 0.002 | 0.006 |  |  |  |  |
| Model 60 - toehold switch sequences | Valeri et al. (18) | 0.135 | - | - | 0.680 | - | - | - | - |
|  | BioAutoML-FAST | 0.180 | 0.051 | 0.226 | 0.484 | - | - | - | - |
| | | ± | ± | ± | ± | | | | |
|  |  | 0.002 | 0.001 | 0.002 | 0.008 |  |  |  |  |

We emphasize that SOTA comparisons are based on reported results from the original studies, which vary in terms of dataset splits, the number of runs, and statistical reporting. Our goal is not to claim absolute superiority, but to demonstrate that a fully automated, standardized pipeline can achieve competitive performance across a wide range of biological sequence tasks.

### Case study 1: Creating a predictor

BioAutoML-FAST generates highly effective predictors for antimalarial peptides, an important target in the effort to combat neglected tropical diseases, as shown in Table 5. Notably, the platform outperforms ESM-2 across most antimalarial peptide metrics while offering greater interpretability by using handcrafted features rather than pLM embeddings. Building on this capability, Figure 2 presents selected results from training an antimalarial predictor, including model metadata, selected descriptors, and feature importance.

**Fig. 2.**
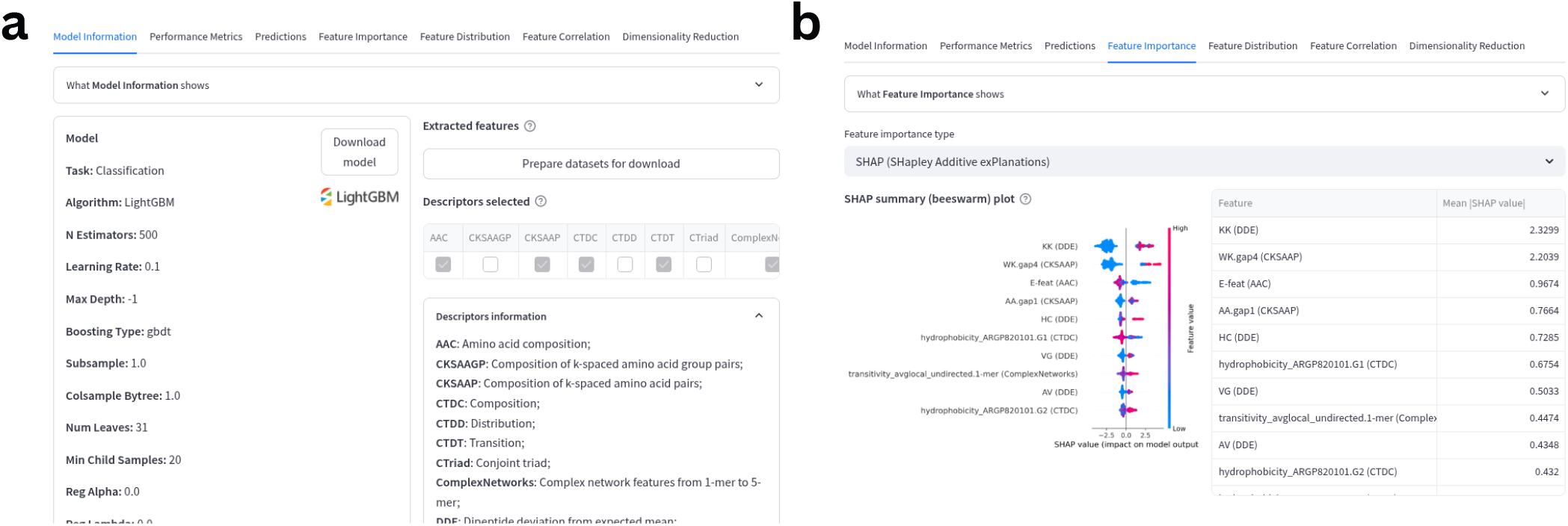
Antimalarial model training and performace. **(a)** Model summary panel displaying the optimal algorithm, hyperparameters, and descriptors identified by the AutoML pipeline. Users can download the trained model object and the corresponding feature matrices for reproducibility. **(b)** Model interpretability using SHAP values to visualize and rank the molecular descriptors with the highest impact on prediction accuracy.

The feature sets (Figure 2a) AAC, CKSAAP, CTDC, and CTDT were automatically selected to better capture the multifaceted physicochemical mechanisms governing antimalarial activity. AAC establishes a baseline by quantifying residue abundance, while CKSAAP complements this by encoding local motifs and spacing patterns critical for secondary structure formation. Furthermore, CTDC and CTDT explicitly model the distributions and transitions of physicochemical properties (e.g., hydrophobicity) that are essential for lipid bilayer disruption. This selection is validated in Figure 2b, which shows that a broad range of the features from those descriptors significantly contribute to the model’s predictive performance.

### Case study 2: Using the model repository to predict unlabeled sequences

To avoid training models from scratch, users can use the Model Repository, which contains 60 predictors for immediate application to novel, unlabeled sequences. The platform excels in non-coding sequence analysis, handling tasks ranging from binary classification to complex 8-class problems. A standout example is Model 55, which robustly distinguishes between diverse RNA biotypes, including miRNA, mRNA, pre-miRNA, rRNA, snoRNA, snRNA, tmRNA, and tRNA. As illustrated in Figure 3, users can view the predicted outcomes for each submitted sequence.

**Fig. 3.**
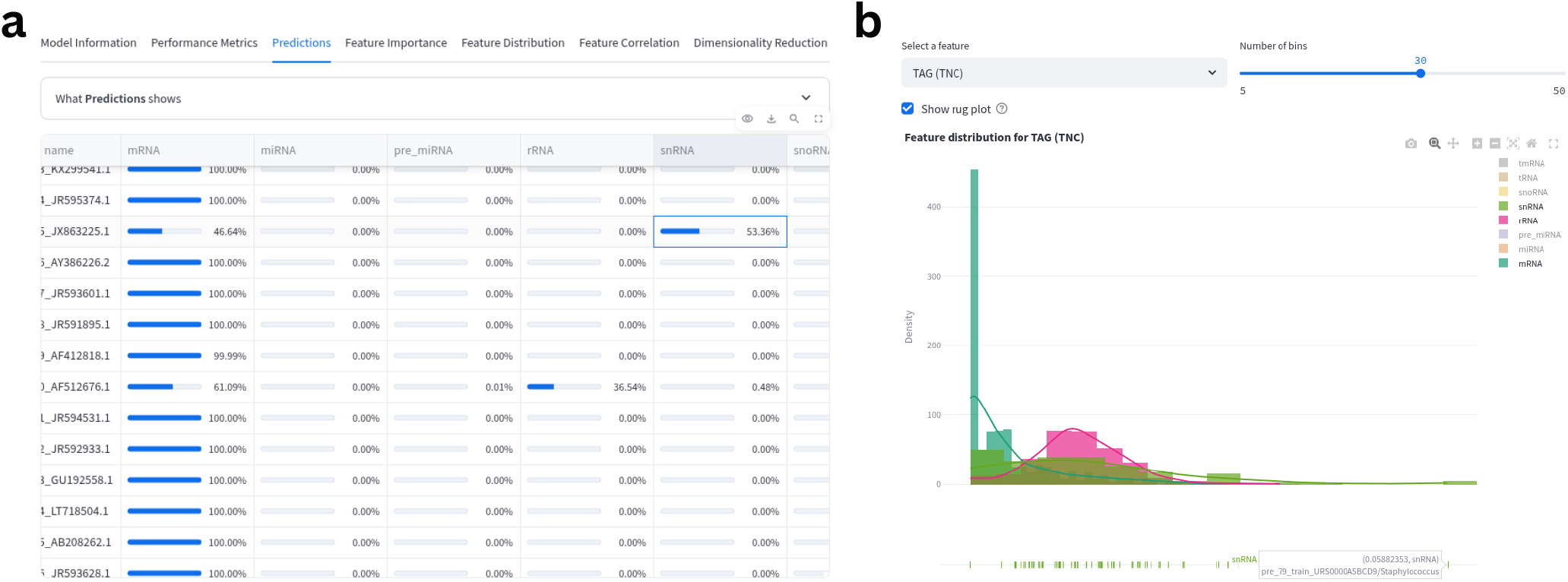
Predicting unlabeled sequences. **(a)** This table summarizes the class probability outputs produced by the Model Repository when a FASTA file containing unlabeled sequences is submitted for prediction. **(b)** Feature distribution of the motif TAG across the classes mRNA, rRNA, and snRNA.

**Fig. 4.**
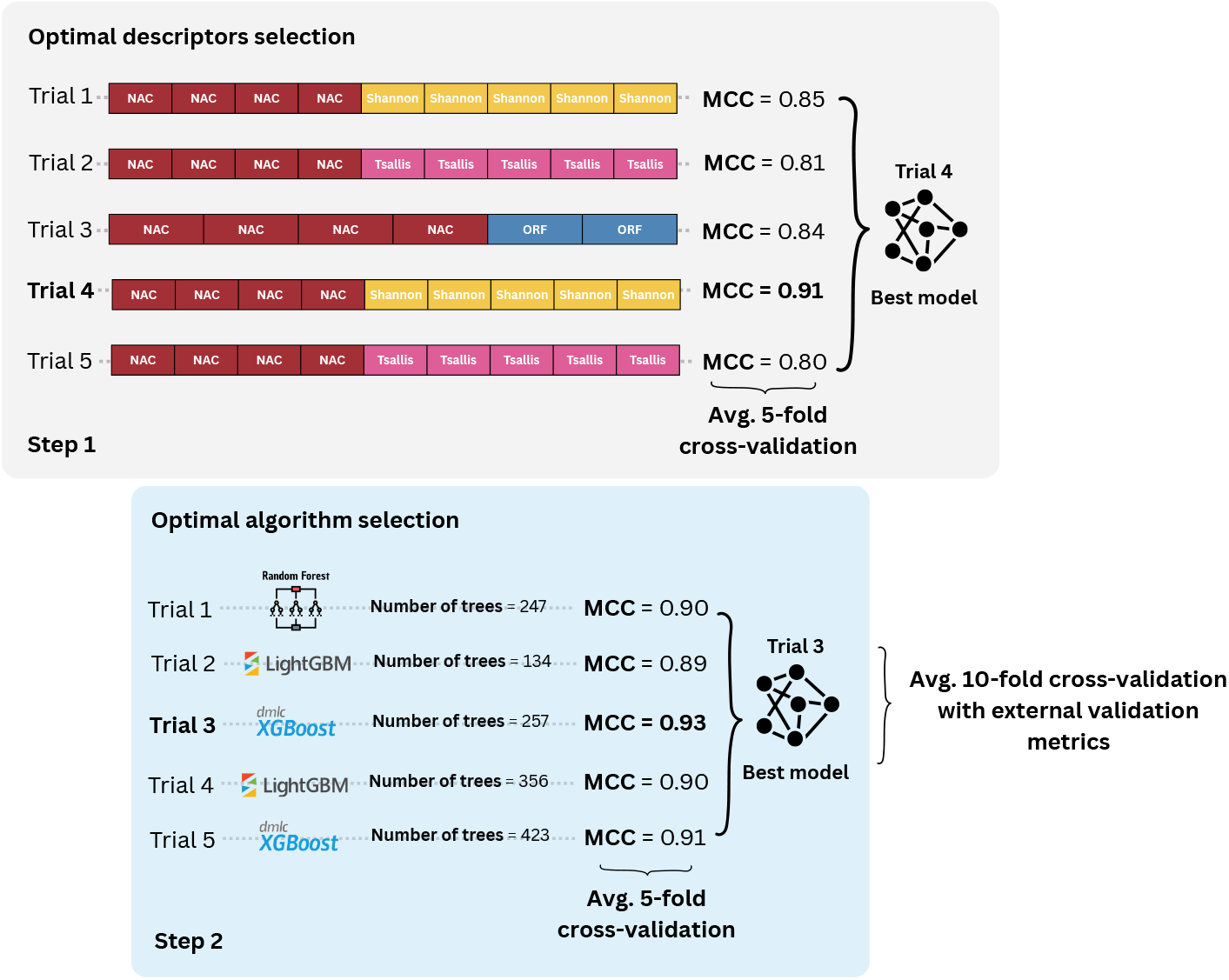
Illustrative example of the 2-step pipeline in BioAutoML-FAST. The framework uses TPE-based Bayesian optimization to identify the optimal predictive model. In step 1, the learning algorithm is fixed (e.g., LightGBM) and the most informative combination of feature descriptors (e.g., NAC, Shannon, Tsallis, ORF) is selected based on mean 5-fold cross-validation performance. In step 2, the selected descriptors are used to jointly optimize the learning algorithm and its hyperparameters, also based on the mean 5-fold cross-validation performance. The best-performing configuration is then forwarded to the final modeling stage for refined 10-fold cross-validation and, when available, external validation on an independent test set.

Even with curated mRNA inputs, the model produces a range of probability scores, reflecting the intrinsic complexity of biological sequences and, in some cases, partial overlap with other RNA classes such as rRNA and snRNA (Figure 3a). This reinforces the need for expert interpretation and for defining application-specific confidence thresholds. Feature-importance analysis showed that the most informative biological signals were Fourier-based sequence-order descriptors, such as coefficient of variation, variance, and kurtosis, as well as gapped *k*-mer statistics, which capture short-range motif-like patterns. Fourier-complex features help distinguish mRNAs through codon-related spectral repetition, whereas kGap features capture conserved nucleotide patterns more typical of structured non-coding RNAs. Partial class overlap may arise when input RNAs are short fragments, such as untranslated regions or incomplete transcripts, that lack strong coding signatures and therefore resemble structured non-coding RNAs or rRNA-derived segments more closely in feature space. To investigate these cases, users can examine feature distributions across classes and assess the behavior of relevant motifs in their dataset. For example, the TAG dispersion shown in Figure 3b, corresponding to the UAG stop codon, displays a strong class-dependent distribution: mRNAs tend to cluster near zero TAG frequency, consistent with stop-codon avoidance in coding regions, whereas non-coding RNAs show broader variation. Examining such distributions can help clarify why certain sequences are difficult to discriminate and where the model is more likely to fail.

## Conclusions

We developed BioAutoML-FAST to democratize access to rigorous ML workflows, addressing the dual challenges of technical complexity and a lack of standardization in biological sequence analysis. By automating the pipeline from feature engineering to deployment, the server enables researchers to perform high-quality analysis without extensive programming expertise. Our evaluation across 60 benchmark datasets confirms that BioAutoML-FAST yields highly competitive performance, matching or outperforming SOTA predictors, including large pLMs like ESM-2, on nearly half the independent set metrics.

Our results show that for tasks with limited data or distinctive physicochemical patterns, classical ML with robust feature engineering can rival deep-learning approaches, while remaining far more computationally efficient and interpretable through descriptors such as *k*-mers, enabling direct feature-importance analysis critical for wet-lab validation. At the same time, our benchmarking exposes systemic gaps in the field, including the scarcity of regression datasets and inconsistent evaluation practices, which BioAutoML-FAST addresses through standardized protocols and a curated repository. Beyond methodology, the platform has significant implications for global accessibility: by lowering technical and computational barriers, it empowers researchers in resource-constrained settings to develop and deploy locally relevant models, fostering decentralized innovation, enabling context-specific biological discovery, and promoting a more equitable and participatory scientific ecosystem.

## Acknowledgements

The authors thank the following people for their valuable suggestions and assistance with testing the platform: Sanchita Kamath, Asaad Ataa, Faith Oni, Jamile Souza, Jana Schor, Matthias Bernt, and Bianca Pessa.

## Data availability

The source code of the platform, along with all datasets used to build the model repository, is available at: https://github.com/Bonidia/BioAutoML-FAST. All trained models can be downloaded directly from the platform.

## Bibliography

1. Claudia Manzoni, Demis A Kia, Jana Vandrovcova, John Hardy, Nicholas W Wood, Patrick A Lewis, and Raffaele Ferrari. Genome, transcriptome and proteome: the rise of omics data and their integration in biomedical sciences. Briefings in bioinformatics, 19 (2):286–302, 2018.

2. Jason A Reuter, Damek V Spacek, and Michael P Snyder. High-throughput sequencing technologies. Molecular cell, 58(4):586–597, 2015.

3. Clelton A Santos, Mariana AB Morais, Fernanda Mandelli, Evandro A Lima, Renan Y Miyamoto, Paula MR Higasi, Evandro A Araujo, Douglas AA Paixão, Joaquim M Junior, Maria L Motta, et al. A metagenomic ‘dark matter’enzyme catalyses oxidative cellulose conversion. Nature, 639(8056):1076–1083, 2025.

4. Stephen F Altschul, Warren Gish, Webb Miller, Eugene W Myers, and David J Lipman. Basic local alignment search tool. Journal of molecular biology, 215(3):403–410, 1990.

5. Sean R Eddy. Hidden markov models. Current opinion in structural biology, 6(3):361–365, 1996.

6. Jaina Mistry, Sara Chuguransky, Lowri Williams, Matloob Qureshi, Gustavo A Salazar, Erik LL Sonnhammer, Silvio CE Tosatto, Lisanna Paladin, Shriya Raj, Lorna J Richardson, et al. Pfam: The protein families database in 2021. Nucleic acids research, 49(D1):D412–D419, 2021.

7. Minoru Kanehisa, Miho Furumichi, Yoko Sato, Yuriko Matsuura, and Mari Ishiguro-Watanabe. Kegg: biological systems database as a model of the real world. Nucleic acids research, 53(D1):D672–D677, 2025.

8. Carsten Kemena and Cedric Notredame. Upcoming challenges for multiple sequence alignment methods in the high-throughput era. Bioinformatics, 25(19):2455–2465, 2009.

9. Tymor Hamamsy, James T Morton, Robert Blackwell, Daniel Berenberg, Nicholas Carriero, Vladimir Gligorijevic, Charlie EM Strauss, Julia Koehler Leman, Kyunghyun Cho, and Richard Bonneau. Protein remote homology detection and structural alignment using deep learning. Nature biotechnology, 42(6):975–985, 2024.

10. Jie Ren, Xin Bai, Yang Young Lu, Kujin Tang, Ying Wang, Gesine Reinert, and Fengzhu Sun. Alignment-free sequence analysis and applications. Annual Review of Biomedical Data Science, 1(1):93–114, 2018.

11. Yanrong Ji, Zhihan Zhou, Han Liu, and Ramana V Davuluri. Dnabert: pre-trained bidirectional encoder representations from transformers model for dna-language in genome. Bioinformatics, 37(15):2112–2120, 2021.

12. Thomas Hayes, Roshan Rao, Halil Akin, Nicholas J Sofroniew, Deniz Oktay, Zeming Lin, Robert Verkuil, Vincent Q Tran, Jonathan Deaton, Marius Wiggert, et al. Simulating 500 million years of evolution with a language model. Science, 387(6736):850–858, 2025.

13. Travers Ching, Daniel S Himmelstein, Brett K Beaulieu-Jones, Alexandr A Kalinin, Brian T Do, Gregory P Way, Enrico Ferrero, Paul-Michael Agapow, Michael Zietz, Michael M Hoffman, et al. Opportunities and obstacles for deep learning in biology and medicine. Journal of the royal society interface, 15(141):20170387, 2018.

14. Valerie Chen, Muyu Yang, Wenbo Cui, Joon Sik Kim, Ameet Talwalkar, and Jian Ma. Applying interpretable machine learning in computational biology—pitfalls, recommendations and opportunities for new developments. Nature methods, 21(8):1454–1461, 2024.

15. Zhen Chen, Pei Zhao, Chen Li, Fuyi Li, Dongxu Xiang, Yong-Zi Chen, Tatsuya Akutsu, Roger J Daly, Geoffrey I Webb, Quanzhi Zhao, et al. ilearnplus: a comprehensive and automated machine-learning platform for nucleic acid and protein sequence analysis, prediction and visualization. Nucleic acids research, 49(10):e60–e60, 2021.

16. Robson P Bonidia, Anderson P Avila Santos, Breno LS de Almeida, Peter F Stadler, Ulisses N da Rocha, Danilo S Sanches, and André CPLF de Carvalho. Bioautoml: automated feature engineering and metalearning to predict noncoding rnas in bacteria. Briefings in Bioinformatics, 23(4):bbac218, 2022.

17. Ioannis Tsamardinos, Paulos Charonyktakis, Georgios Papoutsoglou, Giorgos Borboudakis, Kleanthi Lakiotaki, Jean Claude Zenklusen, Hartmut Juhl, Ekaterini Chatzaki, and Vincenzo Lagani. Just add data: automated predictive modeling for knowledge discovery and feature selection. NPJ precision oncology, 6(1):38, 2022.

18. Jacqueline A Valeri, Luis R Soenksen, Katherine M Collins, Pradeep Ramesh, George Cai, Rani Powers, Nicolaas M Angenent-Mari, Diogo M Camacho, Felix Wong, Timothy K Lu, et al. Bioautomated: an end-to-end automated machine learning tool for explanation and design of biological sequences. Cell systems, 14(6):525–542, 2023.

19. Sabrina Pletzl, Armin Haberl, Tony Ross-Hellauer, and Stefan Thalmann. Reproducible automl: An assessment of research reproducibility of no-code automl tools. 2024.

20. Israel Júnior Borges do Nascimento, Hebatullah Abdulazeem, Lenny Thinagaran Vasanthan, Edson Zangiacomi Martinez, Miriane Lucindo Zucoloto, Lasse Østengaard, Natasha Azzopardi-Muscat, Tomas Zapata, and David Novillo-Ortiz. Barriers and facilitators to utilizing digital health technologies by healthcare professionals. NPJ digital medicine, 6(1): 161, 2023.

21. Denghui Liu, Chi Xu, Wenjun He, Zhimeng Xu, Wenqi Fu, Lei Zhang, Jie Yang, Zhihao Wang, Bing Liu, Guangdun Peng, et al. Autogenome: an automl tool for genomic research. Artificial Intelligence in the Life Sciences, 1:100017, 2021.

22. Haopeng Yu, Fan Li, Bibo Yang, Yiman Qi, Dilek Guneri, Wenqian Chen, Zoë AE Waller, Ke Li, and Yiliang Ding. im-seeker: a webserver for dna i-motifs prediction and scoring via automated machine learning. Nucleic Acids Research, 52(W1):W19–W28, 2024.

23. Raúl Fernández-Díaz, Rodrigo Cossio-Pérez, Clement Agoni, Hoang Thanh Lam, Vanessa Lopez, and Denis C Shields. Autopeptideml: a study on how to build more trustworthy peptide bioactivity predictors. Bioinformatics, 40(9):btae555, 2024.

24. Yungeng Liu, Zan Chen, Yuguang Wang, and Yiqing Shen. Autoproteinengine: A large language model driven agent framework for multimodal automl in protein engineering. In Proceedings of the 31st International Conference on Computational Linguistics: Industry Track, pages 422–430, 2025.

25. Ruite Xiang, Christian Domínguez-Dalmases, Albert Cañellas-Solé, and Victor Guallar. amlprot: an automated machine learning library for protein applications. Bioinformatics, 41(10):btaf543, 2025.

26. Kuo-Chen Chou. Some remarks on protein attribute prediction and pseudo amino acid composition. Journal of theoretical biology, 273(1):236–247, 2011.

27. Bin Liu, Fule Liu, Xiaolong Wang, Junjie Chen, Longyun Fang, and Kuo-Chen Chou. Psein-one: a web server for generating various modes of pseudo components of dna, rna, and protein sequences. Nucleic acids research, 43(W1):W65–W71, 2015.

28. Bin Liu, Fule Liu, Longyun Fang, Xiaolong Wang, and Kuo-Chen Chou. repdna: a python package to generate various modes of feature vectors for dna sequences by incorporating user-defined physicochemical properties and sequence-order effects. Bioinformatics, 31 (8):1307–1309, 2015.

29. Alex T Müller, Gisela Gabernet, Jan A Hiss, and Gisbert Schneider. modlamp: Python for antimicrobial peptides. Bioinformatics, 33(17):2753–2755, 2017.

30. Robson P Bonidia, Douglas S Domingues, Danilo S Sanches, and André CPLF De Carvalho. Mathfeature: feature extraction package for dna, rna and protein sequences based on mathematical descriptors. Briefings in bioinformatics, 23(1):bbab434, 2022.

31. Takuya Akiba, Shotaro Sano, Toshihiko Yanase, Takeru Ohta, and Masanori Koyama. Optuna: A next-generation hyperparameter optimization framework. In Proceedings of the 25th ACM SIGKDD international conference on knowledge discovery & data mining, pages 2623–2631, 2019.

32. Margherita Grandini, Enrico Bagli, and Giorgio Visani. Metrics for multi-class classification: an overview. arXiv preprint 2008.05756, 2020.

33. Gaël Varoquaux and Olivier Colliot. Evaluating machine learning models and their diagnostic value. Machine learning for brain disorders, pages 601–630, 2023.

34. Guido Van Rossum and Fred L. Drake. Python 3 Reference Manual. CreateSpace, Scotts Valley, CA, 2009. ISBN 1441412697.

35. Mohammad Khorasani, Mohamed Abdou, and J Hernández Fernández. Web application development with streamlit. Software Development, 498:507, 2022.

36. Peter JA Cock, Tiago Antao, Jeffrey T Chang, Brad A Chapman, Cymon J Cox, Andrew Dalke, Iddo Friedberg, Thomas Hamelryck, Frank Kauff, Bartek Wilczynski, et al. Biopython: freely available python tools for computational molecular biology and bioinformatics. Bioinformatics, 25(11):1422, 2009.

37. Wes McKinney et al. pandas: a foundational python library for data analysis and statistics. Python for high performance and scientific computing, 14(9):1–9, 2011.

38. John D Hunter. Matplotlib: A 2d graphics environment. Computing in science & engineering, 9(3):90–95, 2007.

39. Plotly Technologies Inc. Collaborative data science, 2015.

40. Michael Waskom, Olga Botvinnik, Drew O’Kane, Paul Hobson, Saulius Lukauskas, David C Gemperline, Tom Augspurger, Yaroslav Halchenko, John B. Cole, Jordi Warmenhoven, Julian de Ruiter, Cameron Pye, Stephan Hoyer, Jake Vanderplas, Santi Villalba, Gero Kunter, Eric Quintero, Pete Bachant, Marcel Martin, Kyle Meyer, Alistair Miles, Yoav Ram, Tal Yarkoni, Mike Lee Williams, Constantine Evans, Clark Fitzgerald, Brian, Chris Fonnesbeck, Antony Lee, and Adel Qalieh. mwaskom/seaborn: v0.8.1 (september 2017), September 2017.

41. Lars Buitinck, Gilles Louppe, Mathieu Blondel, Fabian Pedregosa, Andreas Mueller, Olivier Grisel, Vlad Niculae, Peter Prettenhofer, Alexandre Gramfort, Jaques Grobler, Robert Layton, Jake VanderPlas, Arnaud Joly, Brian Holt, and Gaël Varoquaux. API design for machine learning software: experiences from the scikit-learn project. In ECML PKDD Workshop: Languages for Data Mining and Machine Learning, pages 108–122, 2013.

42. Tianqi Chen and Carlos Guestrin. XGBoost: A scalable tree boosting system. In Proceedings of the 22nd ACM SIGKDD International Conference on Knowledge Discovery and Data Mining, KDD ‘16, pages 785–794, New York, NY, USA, 2016. ACM. ISBN 9781-4503-4232-2. doi: 10.1145/2939672.2939785.

43. Guolin Ke, Qi Meng, Thomas Finley, Taifeng Wang, Wei Chen, Weidong Ma, Qiwei Ye, and Tie-Yan Liu. Lightgbm: A highly efficient gradient boosting decision tree. Advances in neural information processing systems, 30:3146–3154, 2017.

44. Scott M Lundberg and Su-In Lee. A unified approach to interpreting model predictions. Advances in neural information processing systems, 30, 2017.

45. Josiah Carlson. Redis in action. Simon and Schuster, 2013.

46. Laurens van der Maaten and Geoffrey Hinton. Visualizing data using t-sne. Journal of machine learning research, 9(Nov):2579–2605, 2008.

47. Leland McInnes, John Healy, and James Melville. Umap: Uniform manifold approximation and projection for dimension reduction. arXiv preprint 1802.03426, 2018.

48. Ge Liu, Haoyang Zeng, Jonas Mueller, Brandon Carter, Ziheng Wang, Jonas Schilz, Geraldine Horny, Michael E Birnbaum, Stefan Ewert, and David K Gifford. Antibody complementarity determining region design using high-capacity machine learning. Bioinformatics, 36 (7):2126–2133, 2020.

49. Leyi Wei, Chen Zhou, Huangrong Chen, Jiangning Song, and Ran Su. Acpred-fl: a sequence-based predictor using effective feature representation to improve the prediction of anti-cancer peptides. Bioinformatics, 34(23):4007–4016, 2018.

50. Zohre Hajisharifi, Moien Piryaiee, Majid Mohammad Beigi, Mandana Behbahani, and Hassan Mohabatkar. Predicting anticancer peptides with chou s pseudo amino acid composition and investigating their mutagenicity via ames test. Journal of theoretical biology, 341: 34–40, 2014.

51. Piyush Agrawal, Dhruv Bhagat, Manish Mahalwal, Neelam Sharma, and Gajendra PS Raghava. Anticp 2.0: an updated model for predicting anticancer peptides. Briefings in bioinformatics, 22(3), 2021.

52. Patrick Brendan Timmons and Chandralal M Hewage. Ennavia is a novel method which employs neural networks for antiviral and anti-coronavirus activity prediction for therapeutic peptides. Briefings in bioinformatics, 22(6), 2021.

53. Sergio A Pinacho-Castellanos, César R García-Jacas, Michael K Gilson, and Carlos A Brizuela. Alignment-free antimicrobial peptide predictors: improving performance by a thorough analysis of the largest available data set. Journal of Chemical Information and Modeling, 61(6):3141–3157, 2021.

54. Balachandran Manavalan, Shaherin Basith, Tae Hwan Shin, Leyi Wei, and Gwang Lee. mahtpred: a sequence-based meta-predictor for improving the prediction of antihypertensive peptides using effective feature representation. Bioinformatics, 35(16):2757– 2765, 2019.

55. Phasit Charoenkwan, Nalini Schaduangrat, Pietro Lio, Mohammad Ali Moni, Pramote Chumnanpuen, and Watshara Shoombuatong. iamap-scm: a novel computational tool for large-scale identification of antimalarial peptides using estimated propensity scores of dipeptides. ACS omega, 7(45):41082–41095, 2022.

56. Chia-Ru Chung, Ting-Rung Kuo, Li-Ching Wu, Tzong-Yi Lee, and Jorng-Tzong Horng. Characterization and identification of antimicrobial peptides with different functional activities. Briefings in bioinformatics, 21(3):1098–1114, 2020.

57. Xuan Xiao, Pu Wang, Wei-Zhong Lin, Jian-Hua Jia, and Kuo-Chen Chou. iamp-2l: a twolevel multi-label classifier for identifying antimicrobial peptides and their functional types. Analytical biochemistry, 436(2):168–177, 2013.

58. Yuxuan Pang, Lantian Yao, Jingyi Xu, Zhuo Wang, and Tzong-Yi Lee. Integrating transformer and imbalanced multi-label learning to identify antimicrobial peptides and their functional activities. Bioinformatics, 38(24):5368–5374, 2022.

59. Phasit Charoenkwan, Sakawrat Kanthawong, Nalini Schaduangrat, Pietro Li’, Mohammad Ali Moni, and Watshara Shoombuatong. Scmrsa: a new approach for identifying and analyzing anti-mrsa peptides using estimated propensity scores of dipeptides. ACS omega, 7(36):32653–32664, 2022.

60. Lina Zhang, Chengjin Zhang, Rui Gao, Runtao Yang, and Qing Song. Sequence based prediction of antioxidant proteins using a classifier selection strategy. Plos one, 11(9): e0163274, 2016.

61. Ahmad Hassan Butt, Nouman Rasool, and Yaser Daanial Khan. Prediction of antioxidant proteins by incorporating statistical moments based features into chou’s pseaac. Journal of theoretical biology, 473:1–8, 2019.

62. Wei Zhang, Enhua Xia, Ruyu Dai, Wending Tang, Yannan Bin, and Junfeng Xia. Predapp: predicting anti-parasitic peptides with undersampling and ensemble approaches. Interdisciplinary Sciences: Computational Life Sciences, 14(1):258–268, 2022.

63. Phasit Charoenkwan, Chanin Nantasenamat, Md Mehedi Hasan, Balachandran Manavalan, and Watshara Shoombuatong. Bert4bitter: a bidirectional encoder representations from transformers (bert)-based model for improving the prediction of bitter peptides. Bioinformatics, 37(17):2556–2562, 2021.

64. Ruyu Dai, Wei Zhang, Wending Tang, Evelien Wynendaele, Qizhi Zhu, Yannan Bin, Bart De Spiegeleer, and Junfeng Xia. Bbppred: sequence-based prediction of blood-brain barrier peptides with feature representation learning and logistic regression. Journal of Chemical Information and Modeling, 61(1):525–534, 2021.

65. Bin Liu, Jinghao Xu, Xun Lan, Ruifeng Xu, Jiyun Zhou, Xiaolong Wang, and Kuo-Chen Chou. idna-prot|dis: identifying dna-binding proteins by incorporating amino acid distance-pairs and reduced alphabet profile into the general pseudo amino acid composition. PloS one, 9(9):e106691, 2014.

66. Wangchao Lou, Xiaoqing Wang, Fan Chen, Yixiao Chen, Bo Jiang, and Hua Zhang. Sequence based prediction of dna-binding proteins based on hybrid feature selection using random forest and gaussian naive bayes. PloS one, 9(1):e86703, 2014.

67. Bin Liu, Jinghao Xu, Shixi Fan, Ruifeng Xu, Jiyun Zhou, and Xiaolong Wang. Psednapro: Dna-binding protein identification by combining chou’s pseaac and physicochemical distance transformation. Molecular Informatics, 34(1):8–17, 2015.

68. William Stafford Noble, Scott Kuehn, Robert Thurman, Man Yu, and John Stamatoyannopoulos. Predicting the in vivo signature of human gene regulatory sequences. Bioinformatics, 21(suppl_1):i338–i343, 2005.

69. Phasit Charoenkwan, Sakawrat Kanthawong, Chanin Nantasenamat, Md Mehedi Hasan, and Watshara Shoombuatong. idppiv-scm: a sequence-based predictor for identifying and analyzing dipeptidyl peptidase iv (dpp-iv) inhibitory peptides using a scoring card method. Journal of proteome research, 19(10):4125–4136, 2020.

70. Kumardeep Chaudhary, Ritesh Kumar, Sandeep Singh, Abhishek Tuknait, Ankur Gautam, Deepika Mathur, Priya Anand, Grish C Varshney, and Gajendra PS Raghava. A web server and mobile app for computing hemolytic potency of peptides. Scientific reports, 6 (1):22843, 2016.

71. Peng Jiang, Haonan Wu, Jiawei Wei, Fei Sang, Xiao Sun, and Zuhong Lu. Rf-dymhc: detecting the yeast meiotic recombination hotspots and coldspots by random forest model using gapped dinucleotide composition features. Nucleic acids research, 35(suppl_2): W47–W51, 2007.

72. Fatima Khan, Mukhtaj Khan, Nadeem Iqbal, Salman Khan, Dost Muhammad Khan, Abbas Khan, and Dong-Qing Wei. Prediction of recombination spots using novel hybrid feature extraction method via deep learning approach. Frontiers in Genetics, 11:539227, 2020.

73. Siyu Han, Yanchun Liang, Qin Ma, Yangyi Xu, Yu Zhang, Wei Du, Cankun Wang, and Ying Li. Lncfinder: an integrated platform for long non-coding rna identification utilizing sequence intrinsic composition, structural information and physicochemical property. Briefings in bioinformatics, 20(6):2009–2027, 2019.

74. Jun Meng, Qiang Kang, Zheng Chang, and Yushi Luan. Plncrna-hdeep: plant long noncoding rna prediction using hybrid deep learning based on two encoding styles. BMC bioinformatics, 22(Suppl 3):242, 2021.

75. Hao Lv, Zi-Mei Zhang, Shi-Hao Li, Jiu-Xin Tan, Wei Chen, and Hao Lin. Evaluation of different computational methods on 5-methylcytosine sites identification. Briefings in bioinformatics, 21(3):982–995, 2020.

76. Pengmian Feng, Hui Ding, Wei Chen, and Hao Lin. Identifying rna 5-methylcytosine sites via pseudo nucleotide compositions. Molecular BioSystems, 12(11):3307–3311, 2016.

77. Shouzhi Chen, Qing Li, Jianping Zhao, Yannan Bin, and Chunhou Zheng. Neuropred-clq: incorporating deep temporal convolutional networks and multi-head attention mechanism to predict neuropeptides. Briefings in Bioinformatics, 23(5):bbac319, 2022.

78. Yannan Bin, Wei Zhang, Wending Tang, Ruyu Dai, Menglu Li, Qizhi Zhu, and Junfeng Xia. Prediction of neuropeptides from sequence information using ensemble classifier and hybrid features. Journal of proteome research, 19(9):3732–3740, 2020.

79. Yanju Zhang, Sha Yu, Ruopeng Xie, Jiahui Li, André Leier Tatiana T Marquez-Lago, Tatsuya Akutsu, A Ian Smith, Zongyuan Ge, Jiawei Wang, et al. Pengaroo, a combined gradient boosting and ensemble learning framework for predicting non-classical secreted proteins. Bioinformatics, 36(3):704–712, 2020.

80. Lesong Wei, Xiucai Ye, Yuyang Xue, Tetsuya Sakurai, and Leyi Wei. Atse: a peptide toxicity predictor by exploiting structural and evolutionary information based on graph neural network and attention mechanism. Briefings in bioinformatics, 22(5):bbab041, 2021.

81. Phasit Charoenkwan, Chanin Nantasenamat, Md Mehedi Hasan, and Watshara Shoombuatong. Meta-ipvp: a sequence-based meta-predictor for improving the prediction of phage virion proteins using effective feature representation. Journal of Computer-Aided Molecular Design, 34(10):1105–1116, 2020.

82. Mst Shamima Khatun, Md Mehedi Hasan, Watshara Shoombuatong, and Hiroyuki Kurata. Proin-fuse: improved and robust prediction of proinflammatory peptides by fusing of multiple feature representations. Journal of Computer-Aided Molecular Design, 34(12): 1229–1236, 2020.

83. Yiming Zhao, Ningning He, Zhen Chen, and Lei Li. Identification of protein lysine crotonylation sites by a deep learning framework with convolutional neural networks. Ieee Access, 8:14244–14252, 2020.

84. Leyi Wei, Jie Hu, Fuyi Li, Jiangning Song, Ran Su, and Quan Zou. Comparative analysis and prediction of quorum-sensing peptides using feature representation learning and machine learning algorithms. Briefings in Bioinformatics, 21(1):106–119, 2020.

85. Bin Liu, Longyun Fang, Fule Liu, Xiaolong Wang, Junjie Chen, and Kuo-Chen Chou. Identification of real microrna precursors with a pseudo structure status composition approach. PloS one, 10(3):e0121501, 2015.

86. Simon Höllerer, Laetitia Papaxanthos, Anja Cathrin Gumpinger, Katrin Fischer, Christian Beisel, Karsten Borgwardt, Yaakov Benenson, and Markus Jeschek. Large-scale dna-based phenotypic recording and deep learning enable highly accurate sequence-function mapping. Nature communications, 11(1):3551, 2020.

87. Hao Lin, Zhi-Yong Liang, Hua Tang, and Wei Chen. Identifying sigma70 promoters with novel pseudo nucleotide composition. IEEE/ACM transactions on computational biology and bioinformatics, 16(4):1316–1321, 2017.

88. Ranjan Kumar Barman, Anirban Mukhopadhyay, and Santasabuj Das. An improved method for identification of small non-coding rnas in bacteria using support vector machine. Scientific reports, 7(1):46070, 2017.

89. Jacqueline A Valeri, Katherine M Collins, Pradeep Ramesh, Miguel A Alcantar, Bianca A Lepe, Timothy K Lu, and Diogo M Camacho. Sequence-to-function deep learning frameworks for engineered riboregulators. Nature communications, 11(1):5058, 2020.

90. Phasit Charoenkwan, Chanin Nantasenamat, Md Mehedi Hasan, and Watshara Shoombuatong. ittca-hybrid: Improved and robust identification of tumor t cell antigens by utilizing hybrid feature representation. Analytical biochemistry, 599:113747, 2020.

91. Phasit Charoenkwan, Janchai Yana, Chanin Nantasenamat, Md Mehedi Hasan, and Watshara Shoombuatong. iumami-scm: a novel sequence-based predictor for prediction and analysis of umami peptides using a scoring card method with propensity scores of dipeptides. Journal of Chemical Information and Modeling, 60(12):6666–6678, 2020.

92. Junzhe Cai, Ting Wang, Xi Deng, Lin Tang, and Lin Liu. Gm-lncloc: Lncrnas subcellular localization prediction based on graph neural network with meta-learning. BMC genomics, 24(1):52, 2023.

93. Saleh Musleh, Mohammad Tariqul Islam, Rizwan Qureshi, Nehad M Alajez, and Tanvir Alam. Mslp: mrna subcellular localization predictor based on machine learning techniques. BMC bioinformatics, 24(1):109, 2023.

94. Anderson P Avila Santos, Breno LS de Almeida, Robson P Bonidia, Peter F Stadler, Polonca Stefanic, Ines Mandic-Mulec, Ulisses Rocha, Danilo S Sanches, and André CPLF de Carvalho. Biodeepfuse: a hybrid deep learning approach with integrated feature extraction techniques for enhanced non-coding rna classification. RNA biology, 21(1):410–421, 2024.

95. Lezheng Yu, Runyu Jing, Fengjuan Liu, Jiesi Luo, and Yizhou Li. Deepacp: a novel computational approach for accurate identification of anticancer peptides by deep learning algorithm. Molecular Therapy Nucleic Acids, 22:862–870, 2020.

96. Qingwen Li, Wenyang Zhou, Donghua Wang, Sui Wang, and Qingyuan Li. Prediction of anticancer peptides using a low-dimensional feature model. Frontiers in Bioengineering and Biotechnology, 8:892, 2020.

97. Zhenjiao Du, Xingjian Ding, Yixiang Xu, and Yonghui Li. Unidl4biopep: a universal deep learning architecture for binary classification in peptide bioactivity. Briefings in Bioinformatics, 24(3):bbad135, 2023.

98. Phasit Charoenkwan, Wararat Chiangjong, Vannajan Sanghiran Lee, Chanin Nantasenamat, Md Mehedi Hasan, and Watshara Shoombuatong. Improved prediction and characterization of anticancer activities of peptides using a novel flexible scoring card method. Scientific reports, 11(1):3017, 2021.

99. Xuan Xiao, Yu-Tao Shao, Xiang Cheng, and Biljana Stamatovic. iamp-ca2l: a new cnnbilstm-svm classifier based on cellular automata image for identifying antimicrobial peptides and their functional types. Briefings in bioinformatics, 22(6):bbab209, 2021.

100. Luu Ho Thanh Lam, Ngoc Hoang Le, Le Van Tuan, Ho Tran Ban, Truong Nguyen Khanh Hung, Ngan Thi Kim Nguyen, Luong Huu Dang, and Nguyen Quoc Khanh Le. Machine learning model for identifying antioxidant proteins using features calculated from primary sequences. Biology, 9(10):325, 2020.

101. Shahana Yasmin Chowdhury, Swakkhar Shatabda, and Abdollah Dehzangi. idnaprot-es: identification of dna-binding proteins using evolutionary and structural features. Scientific reports, 7(1):14938, 2017.

102. Hong-Liang Li, Yi-He Pang, and Bin Liu. Bioseq-blm: a platform for analyzing dna, rna and protein sequences based on biological language models. Nucleic acids research, 49(22): e129–e129, 2021.

103. Bin Liu, Ren Long, and Kuo-Chen Chou. idhs-el: identifying dnase i hypersensitive sites by fusing three different modes of pseudo nucleotide composition into an ensemble learning framework. Bioinformatics, 32(16):2411–2418, 2016.

